# Effect of cannabidiol on apoptosis and cellular interferon and interferon-stimulated gene responses to the SARS-CoV-2 genes *ORF8*, *ORF10* and *M protein*

**DOI:** 10.1101/2022.01.11.475901

**Authors:** Maria Fernanda Fernandes, John Zewen Chan, Chia Chun Joey Hung, Michelle Victoria Tomczewski, Robin Elaine Duncan

## Abstract

**Aims:** To study effects on cellular innate immune responses to novel genes *ORF8* and *ORF10,* and the more conserved *Membrane protein* (*M protein*) from the Severe acute respiratory syndrome coronavirus 2 (SARS-CoV-2) that causes COVID-19, either alone, or in combination with cannabidiol (CBD).

**Main Methods:** HEK293 cells were transfected with a control plasmid, or plasmids expressing *ORF8*, *ORF10*, or *M protein*, and assayed for cell number and markers of apoptosis at 24 h, and expression of interferon and interferon-stimulated genes at 14 h.

**Key findings:** A significant reduction in cell number, and increase in early and late apoptosis, was found after 24 h in cells where expression of viral genes was combined with 1-2 μM CBD treatment, but not in control-transfected cells treated with CBD, or in cells expressing viral genes but treated only with vehicle. CBD (2 μM) augmented expression of *IFNγ*, *IFNλ1* and *IFNλ2/3*, as well as the 2’-5’-oligoadenylate synthetase (OAS) family members *OAS1*, *OAS2*, *OAS3*, and *OASL,* in cells expressing *ORF8*, *ORF10*, and *M protein*. CBD also augmented expression of these genes in control cells not expressing viral genes, without enhancing apoptosis.

**Significance:** Our results demonstrate a poor ability of HEK293 cells to respond to SARS-CoV-2 genes alone, but suggest an augmented innate anti-viral response to these genes in the presence of CBD. Furthermore, our results indicate that CBD may prime components of the innate immune system, increasing readiness to respond to viral infection without activating apoptosis, and therefore could be studied for potential in prophylaxis.

## 1. Introduction

Coronavirus disease 2019 (COVID-19) is caused by the Severe Acute Respiratory Syndrome Coronavirus 2 (SARS-CoV-2) that was first detected in humans in that year [1]. At the time of writing, the number of cases of COVID-19 is approaching 250 million globally [2], and a number of SARS-CoV-2 variants have emerged and spread between continents [3–8]. Although an effective vaccine is the ultimate goal, efforts to slow the spread, reduce transmission and infectivity, improve health outcomes, and mitigate the most serious health impacts of this disease, will require a multi-faceted approach to reduce the medical, social, and economic burdens of COVID-19. In this regard, the development of effective therapeutics and prophylactics will be key to any effective global health strategy and are urgently needed.

The SARS-CoV-2 genome has been sequenced [9], and found to share significant homology with the genome of SARS-CoV-1, the virus that caused a deadly outbreak of respiratory disease shortly after the turn of the millennium [10]. This homology is fortunate, since prior genomic translational studies, and studies on the cellular function of SARS-CoV-1 viral proteins, have provided some insight into the nature of many of the proteins that function to create the SARS CoV-2 pathogen, and cause COVID-19. However, the SARS-CoV-2 genome has been found to code for an additional novel protein, open reading frame 10 (ORF10) protein, that was not encoded in the SARS-CoV-1 genome, and therefore a function for this protein cannot be inferred from prior work [10]. Studies on this protein have suggested that it is not necessary for virulence or infectivity [11], although sequence analysis indicates that it contains multiple cytotoxic T lymphocyte epitopes [12]. While it is known to be mutated in variants found in humans [13], the function of ORF10 has not yet been elucidated [14].

In addition to ORF10, other proteins encoded by the SARS-CoV-2 genome are yet poorly understood. The ORF8 protein corresponds to two different proteins in SARS-CoV-1, ORF8a and ORF8b, with which it shares only 38.9% and 44.4% sequence identity, respectively, and which differ significantly in protein structure [15]. The role of ORF8 has been suggested to be ‘involvement in host immune evasion’ [15, 16]. However, studies have variably reported that SARS-CoV-2 variants with deletions leading to a deficiency of ORF8 have no difference in infectivity versus wildtype virus [17], or cause milder infections [18], or may combine with additional spike protein mutations to increase transmissibility [19]. Experimental studies on effects of ORF8 in cells also report diverse findings, including the initiation of endoplasmic reticulum stress [20], and evidence of a role in driving the cytokine storm through activation of the interleukin (IL)-17 pathway [21]. With regards to evidence of a role in host immune evasion, studies report inhibitory effects of ORF8 on the induction of Type I interferons (IFN), particularly IFN-*β*[15, 22]. This is notable in the context of COVID-19, since disrupted innate intracellular anti-viral host defenses are specifically implicated in the pathogenesis of this disease [23].

Unlike adaptive immunity, which is mediated by specialized cells of the immune system, essentially all cells are capable of mounting an innate immune response (although the innate immune response functions, in part, to activate adaptive immunity) [24]. The innate immune response can be initiated by cellular entry of viruses or viral components, such as viral RNA or capsid proteins, which are recognized by host pattern recognition receptors that, in turn, trigger signaling cascades leading to the production of host defense molecules including IFNs [24]. However, viruses frequently evolve strategies to disrupt IFN-mediated signaling, and this is reportedly also a function of several non-structural proteins in the SARS-CoV-2 genome [23].

Type I IFN include IFNα and IFNβ, and are among the earliest cytokines produced during the innate immune response following viral infection of cells [25]. While the functions of Type I IFN are complex and can vary throughout an infection, they tend to act initially in the recruitment of immunocytes to promote activation of the acquired host immune response, inhibit proliferation of infected cells, and limit viral replication [25]. Type II IFN, or IFN*γ*, is involved in macrophage and neutrophil activation, and an absence of this factor results in increased virus replication and decreased survival of mice infected with herpes simplex virus type 2 [26]. Type III, or λ-type IFN (IFNλ) are comprised of IFNλ1, and IFNλ2/IFNλ3, which are ∼95% homologous, and IFNλ4 (although expression of this homologue is suppressed at the mRNA or protein level, so it is typically not detected) [27].

While Type III IFN perform similar roles to Type I IFN and were initially thought to be redundant, they are now recognized to be more pro-apoptotic than Type I or Type II IFN [28]. Lambda-type IFN are of significant interest in COVID-19 as a result of evidence showing their greater efficacy at controlling SARS-CoV-2 replication and spread compared to Type I IFN [29], as well as evidence indicating an inverse correlation between Type III IFN levels and severity of COVID-19 [30]. Among the Type III IFN-stimulated genes (ISG) that act as down-stream effectors to induce apoptosis are the 2’-5’-oligoadenylate synthetase (OAS) family members [31, 32]. OAS proteins act as sensors of cytosolic double-stranded RNA produced when viruses replicate, interacting with and activating RNase L after encountering this viral product [33]. RNase L halts viral replication and viral gene translation by cleaving viral protein-encoding RNAs, and also disrupts the host cell transcriptome by degrading cellular rRNAs and tRNAs [33], promoting apoptosis [34, 35]. This strategy can be highly protective in limiting the initiation and spread of an initial infection [36–38]. Although this system is activated in cells infected with SARS-CoV-2, that activation is weak, in contrast to the activation observed in cells infected with other beta-coronaviruses such as SARS-CoV-1 and Middle East Respiratory Syndrome (MERS-CoV) [39]. Pharmacological strategies to increase activation of the OAS-RNase L pathway have thus been suggested as a priority in COVID-19 [40]. This is strongly supported by findings that a polymorphism in a Neanderthal-lineage variant of the *OAS1* gene inherited by some Europeans is associated with higher circulating levels of OAS1 in the non-infected state, and with significant reductions in the risk of COVID-19 susceptibility (odds ratio (OR) = 0.78), hospitalization (OR = 0.61), and ventilation or death (OR = 0.54) following infection [40].

In the current work, we have undertaken studies to examine the effects of expression of *ORF8* and *ORF10* genes, as well as the SARS-CoV-2 structural *Membrane (M) protein*, which is reported to inhibit Type I and III IFN responses [41], on apoptosis and expression of IFNs and down-stream effectors. In addition to examining the effects of expression of these genes alone, we have also investigated effects of combining their expression with cannabidiol (CBD). CBD is the major non-psychotropic phytocannabinoid constituent of *Cannabis sativa* [42], and has been hypothesized as a potential therapeutic in COVID-19 [43, 44]. Evidence from the literature supports that CBD has anti-inflammatory properties [45] and may have a role as a potential protective agent or therapeutic in cells experiencing metabolic distress, such as that associated with viral infection [42, 46]. Based on this, we hypothesized that SARS-CoV-2 genes would be pro-apoptotic, and that CBD would reverse these effects. Instead, we found a potential role for CBD in augmentation of the innate anti-viral host cell response to the viral genes, with evidence of a role for enhanced *IFN*- and *ISG*-induction. While this was initially unexpected, during preparation of the manuscript, data became available demonstrating that CBD inhibits the infection of cells with SARS-CoV-2, as well as replication of the virus after entry into cells, in association with augmented host-cell IFN responses [47]. Our work now shows evidence that CBD augments the anti-viral innate immune response to three distinct viral genes with apparently disparate functions, and also that CBD may prophylactically prime the innate anti-viral response of cells, allowing them to be better prepared to respond to viral infection.

## 2. Materials and methods

### 2.1 Cell culture

HEK293 (human embryonic kidney) cells were grown in Dulbecco’s Modified Eagle’s Medium (DMEM) supplemented with 10% fetal bovine serum (FBS) and 100 U/mL penicillin and 100 mg/mL streptomycin, at 37°C with 5% CO2. Cells were grown to 80% confluence and then routinely subdivided following trypsin digest, and were used at less than 15 passages. The use of HEK293 cells in this study was approved by the University of Waterloo Research Ethics Board (ORE#42425).

### 2.2 Plasmids, transfections, and treatments

Plasmids expressing ORF8 protein (YP_009724396.1) tagged at the C-terminus with 3 x DYKDDDK tag (Ex-NV229-M14), ORF10 protein (YP_009725255.1) tagged at the C-terminus with 3 x hemagglutinin tag (Ex-NV231-M07), and M protein (YP_009724393.1) tagged at the C-terminus with green fluorescent protein (Ex-NV225-M03) were from GeneCopoeia (Rockland, MD, U.S.A). The control plasmid was pCMV-3Tag-3A (pCMV) (Agilent Technologies, Santa Clara, CA, U.S.A.). HEK293 cells were seeded at a density of 1 × 10^4^ cells per well in either 96- or 24-well plates and transfected 24 hours later using JetPRIME (Polyplus Transfection, New York, NY, U.S.A.), according to the manufacturer’s instructions. Briefly, for transfection in a 96-well plate, 0.1 μg of plasmid DNA and 0.25 μL jetPRIME reagent were mixed with 5 μL buffer and incubated for 10 min at room temperature. For transfection in a 24-well plate, 0.5μg of plasmid DNA and 1.25 μL jetPRIME reagent were mixed with 50 μL buffer and incubated for 10 min at room temperature. The incubated solution was diluted in culture medium to a volume of 100 μL (for 96-well plates) or 500 μL (for 24-well plates) and the mixture replaced the culture medium of the cells. Approximately 2-3 h after transfection, cells were treated with either CBD or vehicle (0.1% ethanol) for 24 h. CBD (# ISO60156-1) was purchased from Cedarlane Labs (Burlington, ON, Canada). All work was performed in accordance with a Health Canada approved Cannabis Tracking and Licencing System Research License held by the University of Waterloo (PI: Dr. Robin Duncan).

### 2.3 Crystal violet staining

Relative cell numbers were quantified using the crystal violet staining method, as previously described [48]. Briefly, HEK293 cells were seeded (1 × 10^4^ cells) in 96-well plates and transfected with the respective plasmids after 24 h, then treated a few hours after transfection with either CBD or vehicle for 24 h. Cells were gently washed with 1x phosphate buffered saline (PBS), fixed with a mixture of 10% methanol (v/v), 10% acetic acid (v/v) and stained with crystal violet (Fisher Scientific, Mississauga, Ontario, Canada), then washed and eluted for measurement of absorbance of the samples using a BioTek Synergy H1 Hybrid Multi-Mode Microplate reader at 595 nm.

### 2.4 Apoptosis assay

Early and late apoptotic cells were detected using a Kinetic Apoptosis Kit (#ab129817, Abcam, Toronto, Ontario, Canada), according to the manufacturer’s instructions. Briefly, cells were seeded (1 × 10^4^ cells) in 96-well plates and allowed to adhere for 24 hours, then transfected and treated with either CBD or vehicle for 24 hours, labelled with Polarity Sensitive Indicator of Viability & Apoptosis (pSIVA™), which detects early/ongoing apoptosis, and with Propidium Iodide (PI), which detects cells that are in late apoptosis. Live cells were maintained at 37° C while fluorescence was recorded at 469/525 nm for the detection of pSIVA and at 531/647 nm for the detection of PI. Results are expressed as an index, with the early apoptosis index calculated as pSIVA absorbance at 525 nm/relative cell number per well, and the late apoptosis index calculated as PI absorbance at 647 nm/relative cell number per well.

### 2.5 *IFN* and *ISG* mRNA expression

qPCR analysis was conducted as we have previously described [49]. Cells were grown in 24 well plates and transfected with either pCMV-3Tag-3A, or plasmids expressing *ORF8*, *ORF10*, or *M protein*, and then treated with either 2 μM CBD or vehicle overnight for 14 h, so that analyses were performed prior to measures of effects on cell number and apoptosis markers. Total RNA was isolated using TRIzol^®^ Reagent (1 ml per well) as described by the manufacturer (Invitrogen, Waltham, MA). Quantification of RNA samples was performed using a Nanodrop 2000 Spectrophotometer (Thermo Fisher, Waltham, MA) that was also used to check for A260/280 ratio as an indicator of quality, and 2 µg of RNA was used to synthesize cDNA via oligo(dT) priming using a High-Capacity cDNA Reverse Transcription kit from Applied Biosystems (Waltham, MS, USA). For the real-time PCR assays, cDNA was diluted 1:4 and 1 µl was added to a master mix with 9 µl of PerfeCTa SYBR^®^ Green supermix (Quanta Bio, Beverly, MA), 0.5 µl forward and reverse primers (25 µM each) for the targeted gene (please see Table 1 for primer sequences), and 3 µl of ddH_2_0. The cycling conditions for all genes were as follows: 1 cycle of 95°C for 2 min, followed by 49 cycles of 95°C for 10 s, then 60°C for 20 s. Relative expression of the targeted gene was calculated using the *^ΔΔ^*Ct method with the Ct values normalized to glyceraldehyde 3-phosphate dehydrogenase (*GAPDH*).

**Table 1:**
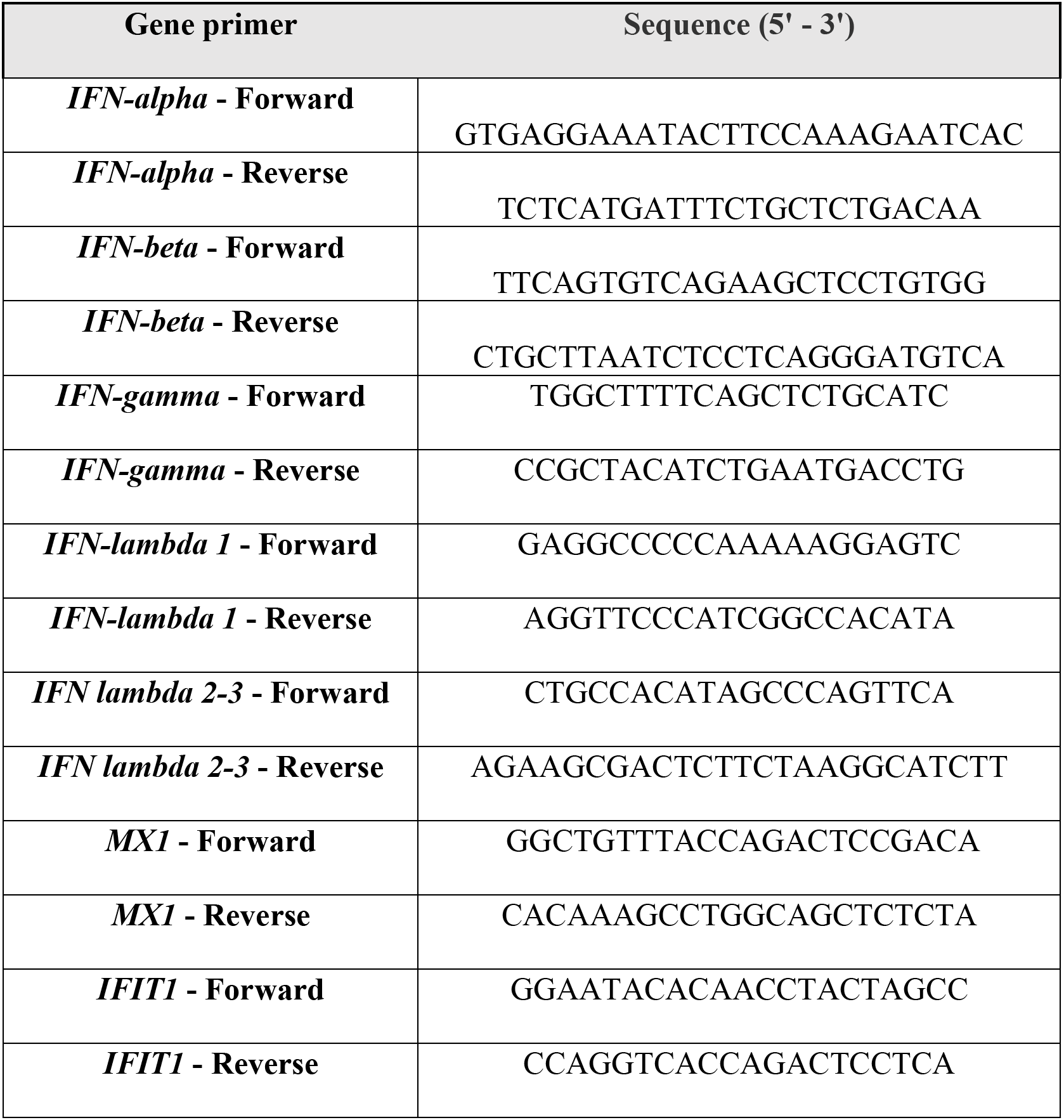
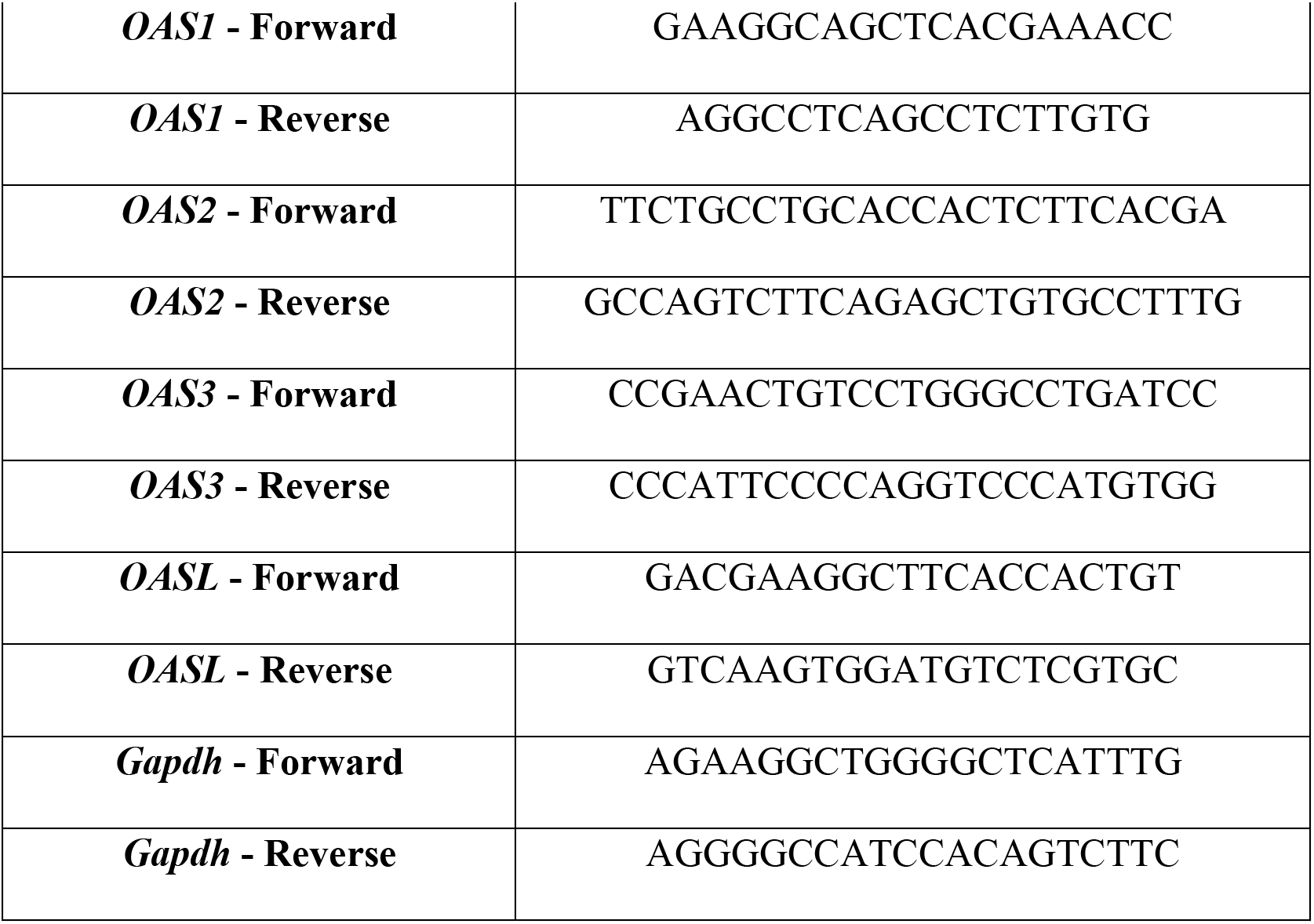
Primer sequences

### 2.6 Statistical analyses

Non-linear regression was performed on data generated from the concentration-dependent effects of CBD on cell number in cells transfected with control and viral gene expression plasmids and used to determine IC50 values for CBD in combination with each viral gene. Simple linear regression was performed to determine if the slopes were significantly non-zero. Two-way analysis of variance (ANOVA) followed by Tukey’s post-hoc test for multiple comparisons was performed to compare early and late apoptosis indexes, and gene expression levels, among cells transfected with control and viral gene-expression plasmids, with and without various concentrations of CBD. Analyses were performed using Prism GraphPad 9 software. Data shown are means ± S.E.M.; n-values denote the number of biological replicates derived from, at a minimum, different passages of cells. Where technical replicates were performed within experiments, these were averaged to derive single values reported as biological replicates.

## 3. Results

### 3.1 Relative cell numbers

A concentration response curve was generated by treating cells transfected with the control plasmid (pCMV) or plasmids expressing viral genes with vehicle (0.1% EtOH (*i.e.* 0 μM CBD)), or with increasing concentrations of CBD (Fig. 1A). The range of concentrations tested was based on pharmacologically achievable blood concentrations observed in human pharmacokinetic studies [50]. The slopes of lines generated from concentration-responses to CBD in cells expressing viral genes were significantly non-zero, indicating a significant relationship between increasing dose of CBD and relative cell number, while the slope of the line for pCMV was not significantly non-zero. IC50 values for CBD concentrations were 0.89 μM for cells expressing *ORF8*, 0.91 μM for cells expressing *ORF10*, 0.99 μM for cells expressing *M protein*, and 7.24 μM for cells transfected with pCMV. At a treatment level of 2 μM CBD, relative cell numbers in wells transfected with viral genes were reduced by ∼55-80% (P<0.0001) relative to cell numbers in wells transfected with viral genes but not treated with CBD or relative to wells transfected with control plasmid and treated with or without 2 μM CBD, among which there were no significant differences.

**Fig. 1.**
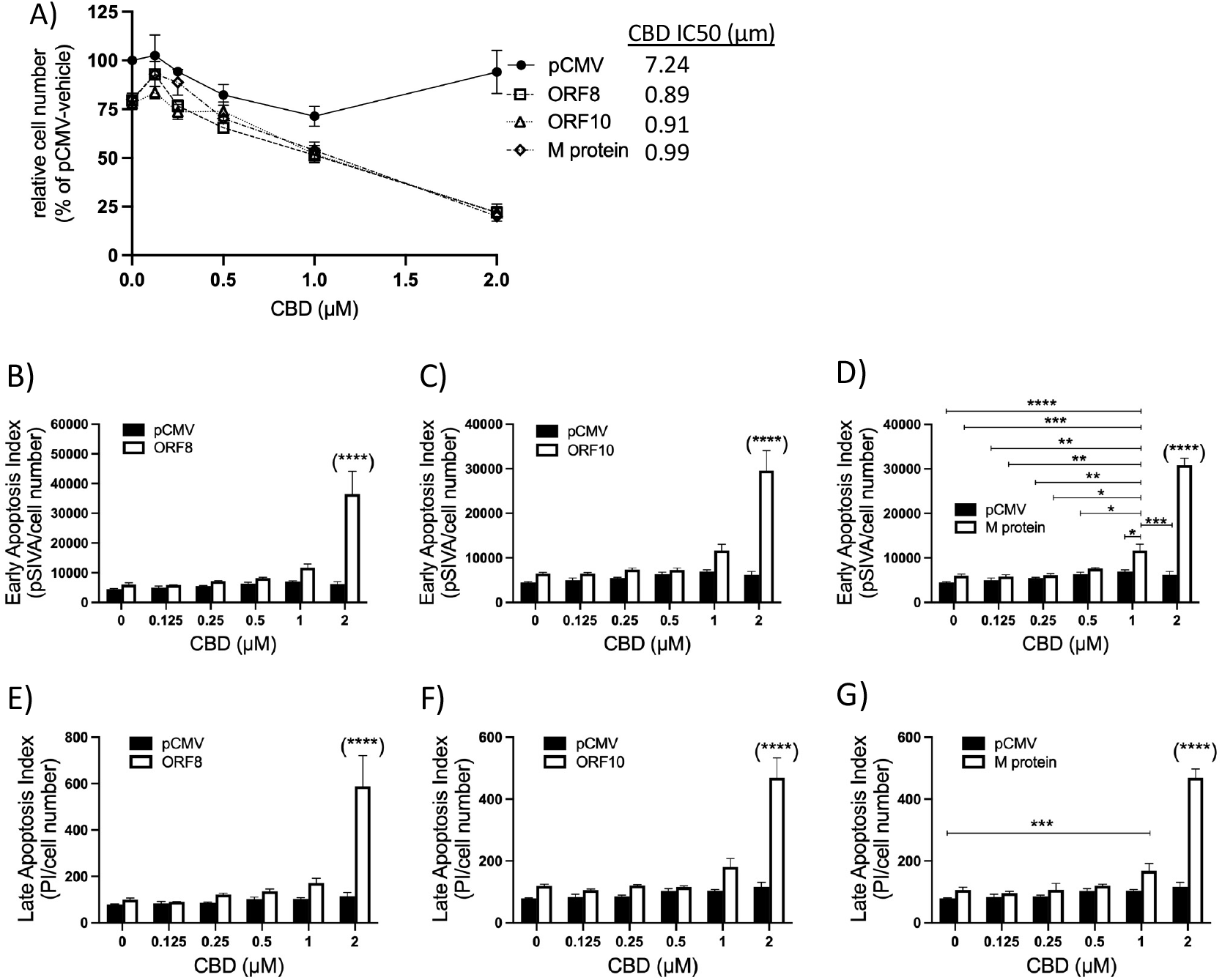
Effect of *ORF8*, *ORF10*, or *M protein* expression, with and without CBD treatment, on HEK293 cell number and apoptosis indexes. (A) Dose-dependent effects of CBD on the relative number of cells per well 24 h after transfection with control plasmid (pCMV), or plasmids expressing *ORF8*, *ORF10*, or *M protein* (n=3-12). IC50 values for CBD concentration in combination with each group are shown. (B-D) Dose-response effect to CBD on the early apoptosis index in HEK293 cells expressing pCMV or viral genes at 24 h. (E-G) Dose-response effect to CBD on the late apoptosis index in HEK293 cells transfected with control or viral plasmids. Apoptotic indexes were calculated by dividing the relative absorption of the respective marker by the number of cells per well. Apoptosis data were analyzed by 2-way ANOVA with Tukey’s post-hoc test, n=3-9. Differences among groups are as indicated, *P<0.05, **P<0.01, ***P<0.001, ****P<0.0001, where ^(****)^ denotes a significant difference (P<0.0001) between cells treated with 2 μM CBD and transfected with a viral gene-encoding plasmid, and all other groups.

### 3.2 Early and late apoptosis

Differences in cell number can result from changes in cell proliferation, or cell death (*i.e.* apoptosis or necrosis), or both. An initial assessment for changes in cell proliferation indicated no significant effect (*data not shown*), and therefore we focused our studies on apoptosis. A concentration-dependent effect of CBD on the activation of an early marker of apoptosis (pSIVA), and on incorporation of a late marker of apoptosis (PI), was evident in cells expressing *ORF8*, *ORF10*, and *M protein*, but this was not observed in cells transfected only with the control plasmid (Figs. 1B-G). Specific analyses comparing cells transfected with the control vector or plasmids expressing viral genes and treated with increasing levels of CBD demonstrate important effects. First, this analysis shows that CBD alone, even at the highest concentration tested, does not significantly increase markers of apoptosis in control cells. Additionally, it demonstrates that expression of the viral genes *ORF8*, *0RF10*, or *M protein* with vehicle alone (*i.e.* 0 μM CBD) also does not significantly increase either early or late apoptosis relative to control cells, indicating a poor ability of cells to detect and respond to the presence of these viral transcripts or proteins in the absence of CBD. Interestingly, however, both early and late apoptosis indexes were significantly elevated in cells expressing any of the viral genes when also treated with 2 μM CBD, relative to all other groups. In cells expressing *ORF8*, early and late apoptosis indexes were both increased by over 6-fold in cells treated with 2 μM CBD compared to indexes in vehicle alone (Fig. 1B, E). In cells expressing *ORF10* (Fig. 1C, F), early and late apoptosis indexes were increased ∼4.7- and ∼4.0-fold, respectively, by 2 μM CBD versus vehicle. In cells expressing *M protein* (Fig. 1D, G), early and late apoptosis indexes were increased by ∼5.6- and ∼4.7-fold in cells expressing *M protein* and treated with 2 μM CBD. In addition, significant effects of 1 μM CBD were also evident on cells expressing *M protein* (Fig. 1D, G). This concentration generated a significantly elevated late apoptosis index relative to vehicle-treated control cells, and significantly greater early apoptosis indexes relative to most other *M protein*-transfected cells at the same or lower levels of CBD treatment, and all other control-transfected cells treated with or without CBD (Fig. 1D, G).

### 3.3 Expression of *IFN* genes

Expression of *IFNα* and *IFNβ* was not significantly altered by *ORF8*, *ORF10*, or *M protein*, either with or without 2 μM CBD (Fig. 2A-F). However, transfection of these viral genes significantly increased the expression of *IFNγ*, and this was augmented by 2 μM CBD (Fig. 3A-C). In the absence of CBD, transfection of cells with *ORF8*, *ORF10*, or *M protein* caused a significant 16- to 29-fold increase in expression of *IFNγ* relative to vehicle-treated control cells, and this effect was augmented by treatment with 2 μM CBD, further increasing *IFNγ* expression (Fig. 3A-C). Interestingly, however, cells transfected with *ORF8* (in the absence of CBD) did not have higher expression of *IFNλ1* or *IFNλ2/3* than controls, although treatment of cells with 2 μM CBD caused a significant induction of *IFNλ1* and *IFNλ2/3* by *ORF8* (Fig. 3D, G). These genes were induced without CBD co-treatment when cells were transfected with *ORF10* (by 9.6-fold and 2.4-fold) (Fig. 3E, H) or *M protein* (by 4.1-fold, for both genes) (Fig. 3F, I) and 2 μM CBD strongly augmented the induction of both *IFNλ1* and *IFNλ2/3* that occurred when *ORF10* or *M protein* were transfected, by a further 3.8- to 11.2-fold (Fig. 3 E, F, H, I). Although relative cell number and apoptosis measures were not significantly affected by 2 μM CBD in pCMV-transfected cells, this treatment caused an ∼5-fold increase in expression of *IFNγ* in pCMV-transfected control cells compared to pCMV-controls cells treated only with vehicle. Similarly, *IFNλ1* and *IFNλ2/3* were increased in pCMV-transfected control cells treated with 2 μM CBD by 3-fold and 7-fold, respectively.

**Fig. 2.**
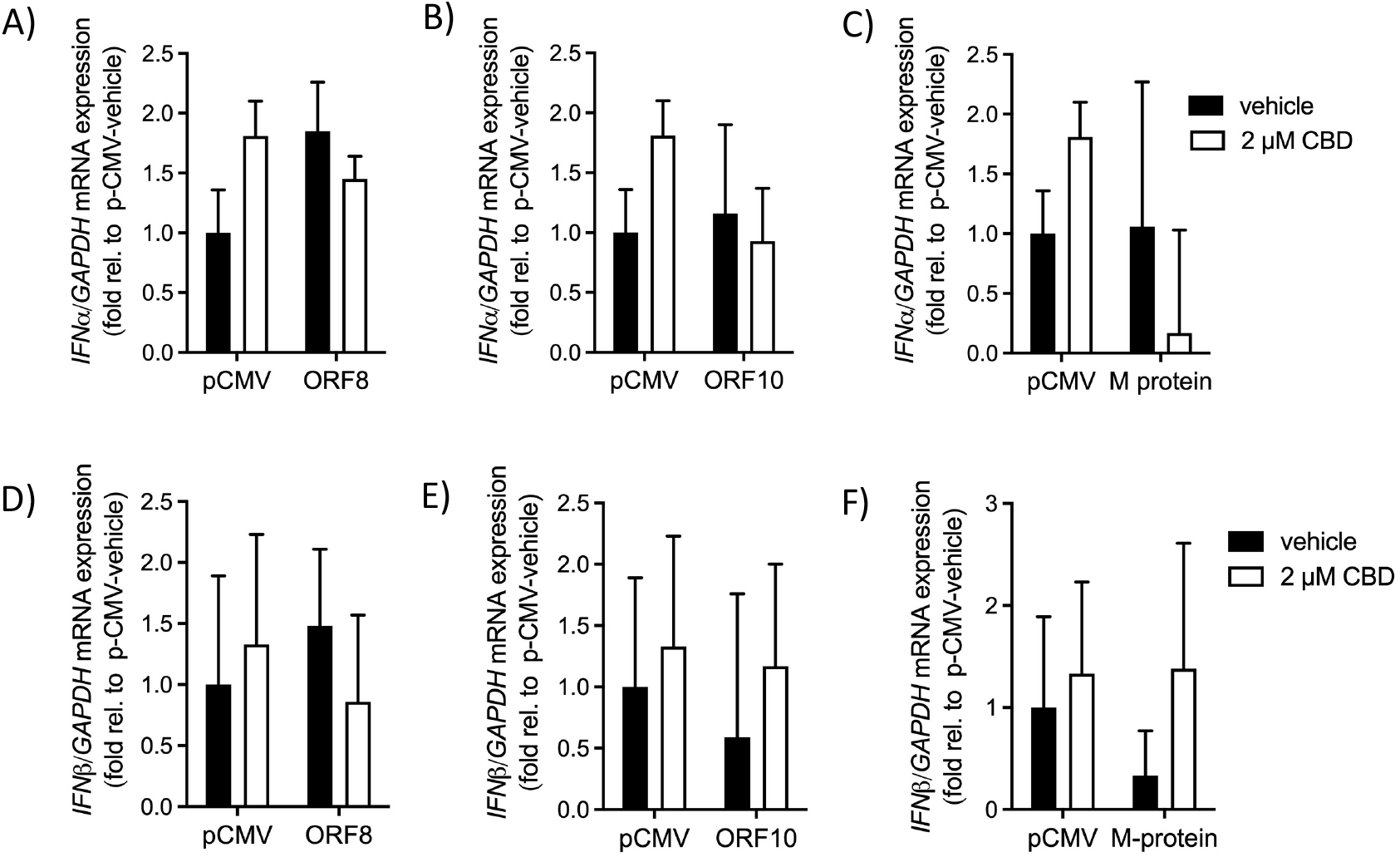
Effect of *ORF8*, *ORF10*, or *M protein*, with and without CBD, on gene expression of Type I *IFN*. Expression of *IFNα* (A-C) and *IFNβ* (D-F) in cells transfected with control plasmid (pCMV), or plasmids expressing *ORF8*, *ORF10*, or *M protein*, and treated with vehicle control (0.1% ethanol) or 2 µm CBD for 14 h. Data are means ± SEM (n=5).

**Fig. 3.**
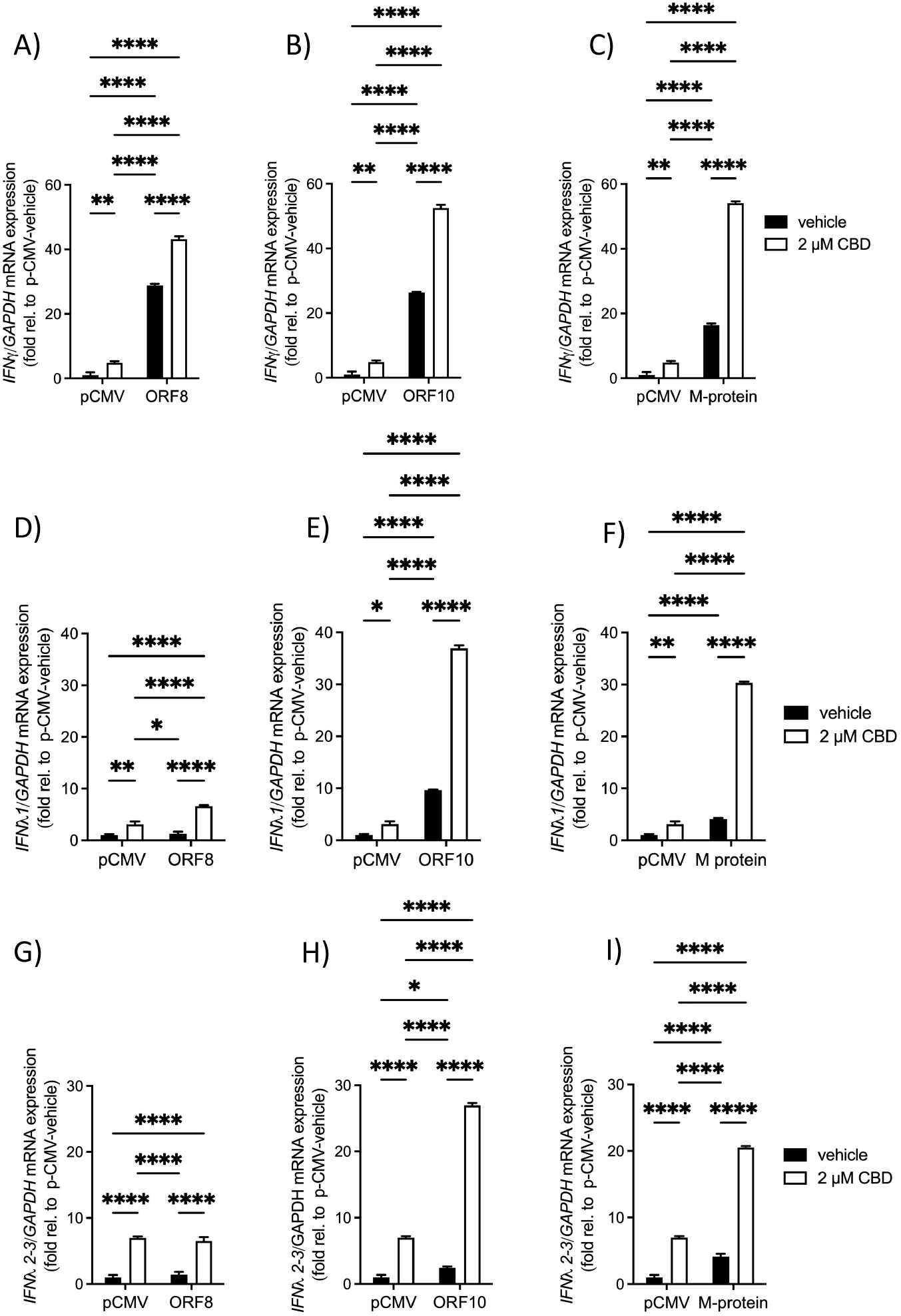
Effect of *ORF8*, *ORF10*, or *M protein*, with and without CBD, on gene expression of Type II and III *IFN*. Expression of *IFNγ* (A-C), *IFNλ1* (D-F), and *IFNλ2/3* (G-I), in cells transfected with control plasmid (pCMV), *ORF8*, *ORF10*, or *M protein*, and treated with vehicle control (0.1% ethanol) or 2 µm CBD (n=5) for 14 hours. Data are means ± SEM, *P<0.05, **P<0.01, ***P<0.001, and ****P<0.0001.

### 3.4 Expression of ISG

Expression of the ISGs *IFIT1* and *MX1* was not significantly altered by treatment with 2 μM CBD, or by expression of the SARS-CoV-2 genes *ORF8*, *ORF10*, and *M protein*, either alone, or in combination (Fig. 4A-F). However, significant effects were observed when *OAS* family genes were analyzed. Surprisingly, transfection of *ORF8*, *ORF10*, and *M protein* did not significantly induce expression of *OAS1*, *OAS2*, or *OAS3* relative to cells transfected with pCMV in the absence of CBD (Fig. 5A-I). This indicates that these cells may have a poor ability to recognize and respond to these viral genes through innate immune system activation involving the OAS family. Only *OASL* was significantly induced by *ORF8* (by 17.9-fold), *ORF10* (by 4.9-fold), and *M protein* (by 18.8-fold), in the absence of CBD (Fig. 5J-L). When 2 μM CBD was added to cells transfected only with the control plasmid, expression of *OAS2, OAS3,* and *OASL* increased significantly (from 5.7 to 7.8-fold). Addition of 2 μM CBD to cells transfected with *ORF8*, *ORF10*, or *M protein*, augmented the expression of all *OAS* family genes relative to the corresponding vehicle-treated cells, with the additional induction caused by CBD ranging from 3.1- to 22.9-fold.

**Fig. 4.**
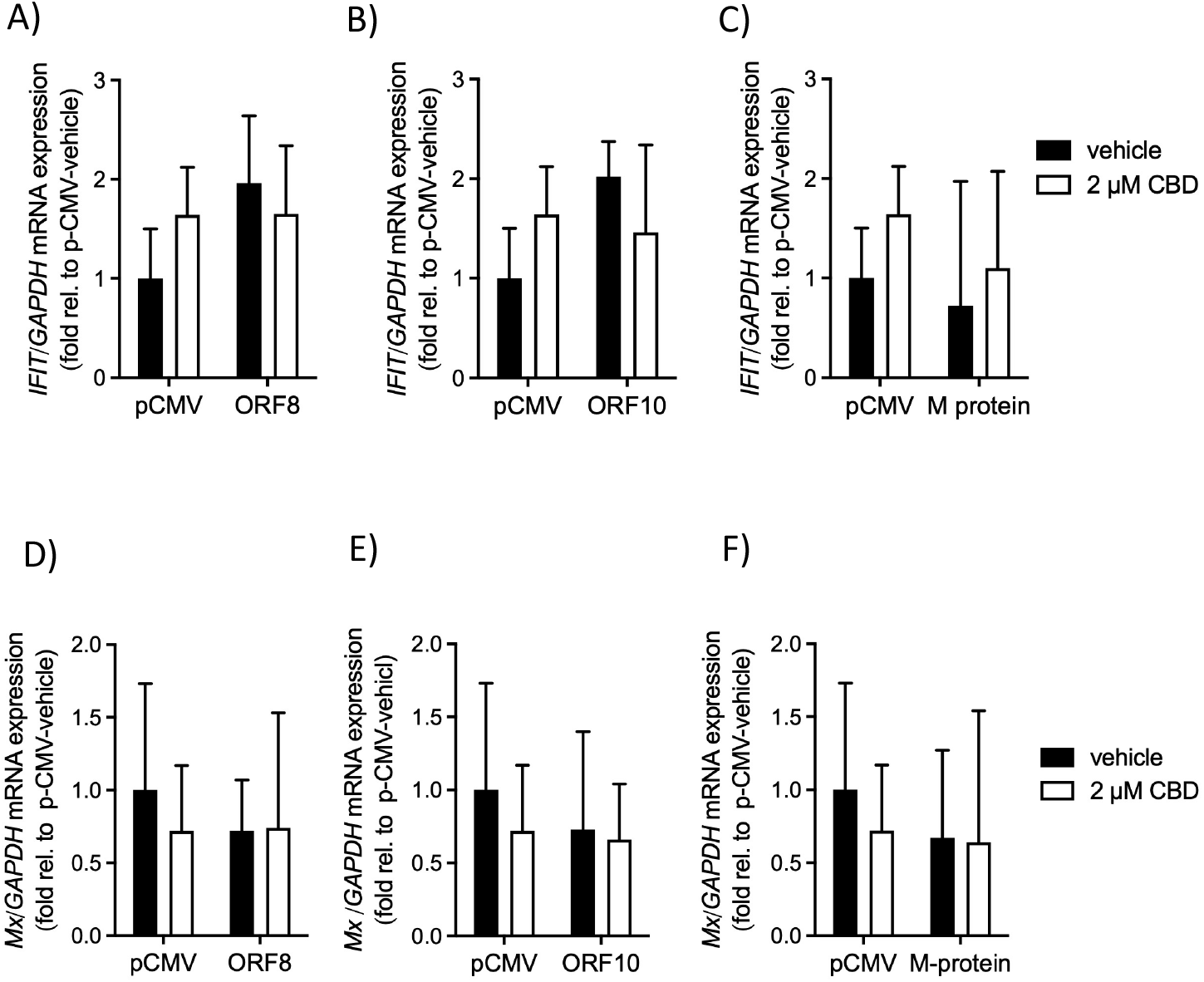
Effect of *ORF8*, *ORF10*, or *M protein*, with and without CBD, on gene expression of *MX1* and *IFIT1*. Expression of *MX1* (A-C) or *IFIT1* (D-F) in cells transfected with control plasmid (pCMV), *ORF8*, *ORF10*, or *M protein*, and treated with vehicle control (0.1% ethanol) or 2 µm CBD for 14 h (n=5). Data are means ± SEM.

**Fig. 5.**
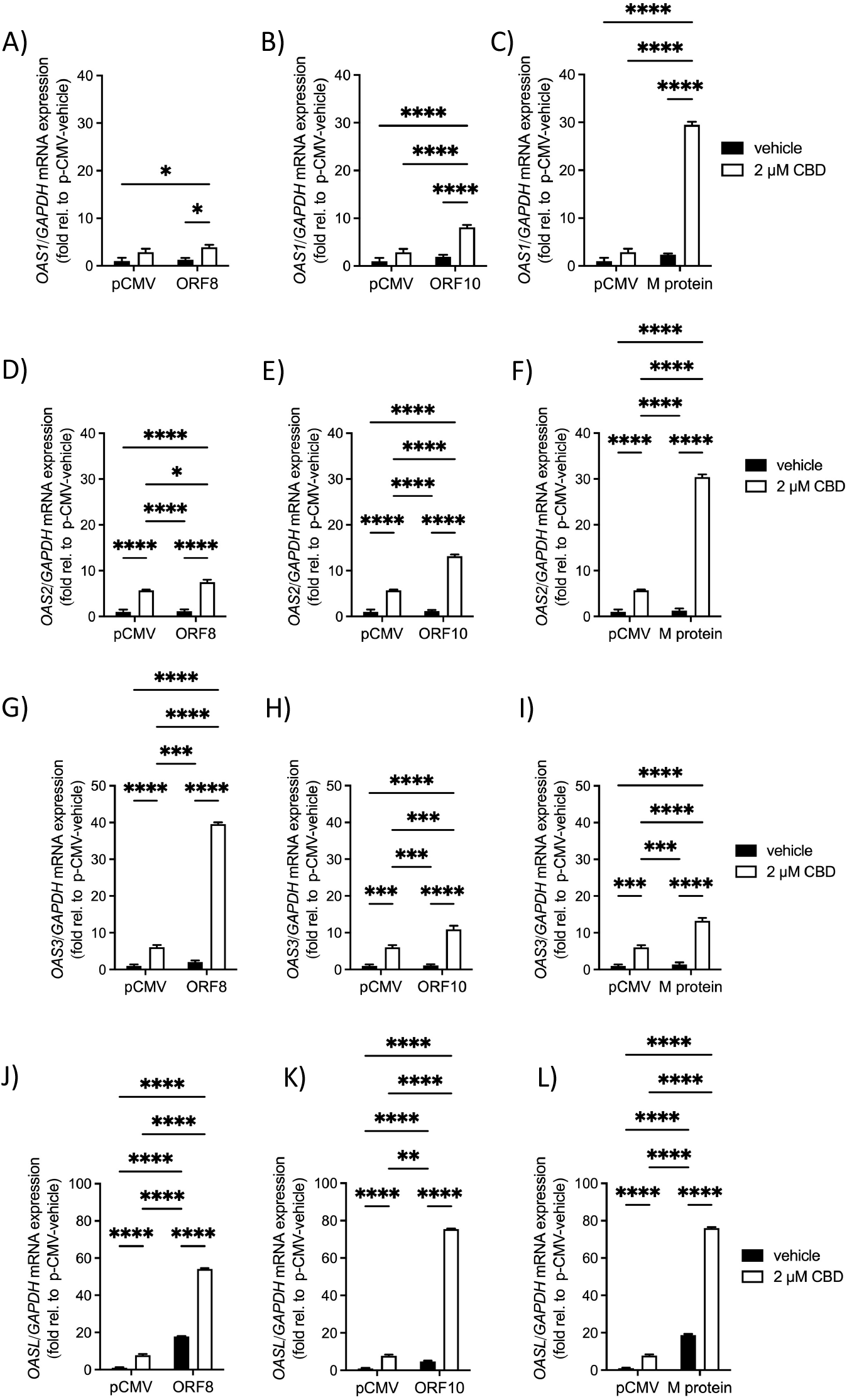
Effect of *ORF8*, *ORF10*, or *M protein*, with and without CBD, on gene expression of *OAS* family members. Expression of *OAS1* (A-C), *OAS2* (D-F), *OAS3* (G-I) and *OASL* (J-L) in cells transfected with control plasmid (pCMV), *ORF8*, *ORF10*, or *M protein*, and treated with vehicle control (0.1% ethanol) or 2 µm CBD for 14 h (n=5). Data are means ± SEM. *P<0.05, ***P<0.001, ****P<0.0001.

## 4. Discussion

The infectious dose of SARS-CoV-2 required to cause disease in 50% of people exposed has been estimated to be 280 virions [51]. Infection with any virus, including SARS-CoV-2, does not initially cause symptoms. At infection, a small number of virus particles enter cells and ‘hijack’ the cellular machinery to replicate, releasing more infectious particles that amplify the titre. Thus, during peak infection, an individual may have 10^9^ to 10^11^ virions in their cells and bodily fluids, which can cause symptomatic disease [52, 53]. Factors that prevent viral replication are of significant interest in the COVID-19 pandemic, since they are protective both for individuals and populations. In a host, replication is needed in order for an initial infectious dose to spread within the body, producing symptomatic disease, although asymptomatic SARS-CoV-2 carriers have been reported [54]. Host replication is also needed to produce a sufficient concentration of viral particles for an individual to become infectious to others in a population [54]. Within a population, widespread replication leads to mutations and the generation of novel variants, which can alter the infectivity and virulence of a virus, and potentially reduce the protective efficacy of vaccines [55].

To redirect the cell’s replicative machinery towards viral production, viruses typically encode proteins that can deregulate cell cycle checkpoints. In coronaviruses, including SARS-CoV-1, the nucleocapsid protein inhibits cell cycle progression and cell proliferation by inhibiting activity of cyclin/cyclin-dependent kinase (CDK) complexes [56]. We observed a concentration-dependent decrease in the number of cells per well when cells were transfected with plasmids expressing *ORF8*, *ORF10*, or *M protein* and treated with CBD, but not when cells were transfected only with the control plasmid. Although we first tested whether expression of *ORF8*, *ORF10* or *M protein* in cells treated CBD would modulate cell proliferation, we did not find any significant differences among groups (*data not shown*). We therefore focused our investigation on a role for these viral genes in modulating apoptosis, which occurs when cells are infected with pathogenic viruses, including SARS-CoV-1 [57] and MERS-CoV [58].

Apoptosis occurs as an outcome of an innate immune response of the cell to viral infection that serves to prevent viral replication and consequently virus spreading and mutation [59]. Cells undergo apoptosis to interrupt the production and release of progeny virus, resulting in early elimination of both the virus and infected cells [60, 61], which may result in the absence of disease, or a milder course of disease, as well as a situation where viral transmission is also prevented or reduced. The induction of apoptosis shortly after exogenous viral genes enter a cell prevents viral genome replication. It is therefore particularly protective against the development of new viral variants, which may potentially arise even in immunized people who can, in some cases, become infected and spread the virus despite vaccination [62].

Interestingly, we found that expression of the SARS-CoV-2 genes *ORF8*, *ORF10*, and *M protein* alone did not significantly induce apoptosis. This is consistent with studies of patients with COVID-19 where the induction of apoptosis was lacking in nasopharyngeal samples [63]. While CBD did not increase apoptosis in control cells, treatment of cells expressing viral genes with a pharmacological dose of CBD significantly augmented the induction of both early and late apoptosis. This finding suggests that CBD may help limit an initial infection by promoting removal of infected cells, thereby limiting the spread, and therefore also likely raising the necessary infectious titre. This is supported by evidence from users of Epidiolex®, a high-dose pharmaceutical CBD licensed in the United States for use in the treatment of rare types of epilepsy in adults and children [47]. In that study, patients prescribed high-dose CBD had an approximate 10-fold lower risk of testing positive for SARS-CoV-2, even when matched by demographics, recorded diagnoses, and other medications. In those with use of any cannabinoid in their medical record, the positivity rate for SARS-CoV-2 was over 40% lower [47]. Taken together with our findings, this suggests that CBD may provide a prophylactic effect against the risk of contracting SARS-CoV-2 and developing COVID-19 by increasing the initial apoptotic response to viral genes.

We investigated the regulation of *IFN* and *ISG* as a potential mechanism underlying this effect. Prior work has indicated that the SARS-CoV-2 virus can counteract host innate anti-viral responses, resulting in suppression of IFN-mediated responses [23]. Thus, factors that can counteract this are of particular interest. We hypothesized that augmented induction of *IFN* and *ISG* could play a role in the enhanced apoptosis observed in cells expressing viral genes and treated with CBD. Interferons are a family of inducible cytokines with pleiotropic biological effects [64], induced at different time points following infection [25], which help to regulate the innate, intracellular, anti-viral host defense [65]. Type I IFNs tend to slow down proliferation and regulate cell survival, while Type II IFNs also regulate cell survival and proliferation, and Type III IFNs induce cell apoptosis, more so than Types I or II [66]. Inadequate induction of IFNs, and especially lambda-type interferons, has been identified as a factor in SARS-CoV-2 infection leading to more severe disease [67]. The IFN λ family are important inducers of the anti-viral immune response at mucosal surfaces [68], and people with a greater IFN λ induction tend to have less viral inflammation, and may not even develop disease [67].

The lack of induction of Type I *IFN* by either viral gene expression or CBD, suggests that these *IFN* were not involved in the pro-apoptotic response observed. In all comparisons, however, Type II and Type III *IFN* were significantly induced by a combination of viral genes and CBD relative to cells expressing only the viral genes without CBD, and in almost all comparisons, also relative to control cells treated with or without CBD. This was similar to observed effects on early- and late-stage apoptosis, where cells expressing viral genes in combination with 2 μM CBD were many fold more effective at inducing apoptosis markers than cells expressing either the viral genes alone, or control plasmid with or without CBD. Although this association between the induction of Type II and III *IFN* and the induction of early- and late-apoptosis is only correlative, it may suggest a possible role for these IFN in mediating observed outcomes.

Analysis of downstream effectors indicated that involvement of *MX1* or *IFIT1* genes was unlikely, although it should be noted that the time course, involving measurements of gene expression preceding apoptosis, may not have captured changes in genes that are typically induced later in the innate immune response [25]. Conversely, *OAS1*, *OAS2*, *OAS3*, and *OASL* family members, which were all significantly elevated in cells expressing viral genes and treated with 2 μM CBD compared to vehicle, were likely factors. Surprisingly, however, expression of *ORF8*, *ORF10*, or *M protein* without CBD was insufficient to induce *OAS1, OAS2, or OAS3* relative to control-transfected cells, in agreement with reports that an inadequate innate immune response of cells to SARS-CoV-2 may be a factor in the pathology of this virus [23]. Of particular note is the finding that control-transfected cells treated with 2 μM CBD expressed significantly higher levels of *IFNγ*, *IFNλ1, IFNλ2/3,* and *OAS2, OAS3, and OASL,* in comparison with control-transfected cells treated only with vehicle, since CBD did not augment apoptosis or significantly reduce cell numbers in these groups. This raises the intriguing possibility that CBD may prime the innate immune system of cells under normal, non-pathological conditions, by raising basal expression of effectors, so that they are better able to recognize and respond to the presence of viral material, upon infection.

Our finding that CBD regulates OAS family gene expression is particularly interesting, given the role of these enzymes as powerful mediators of virus-associated apoptosis [69–72]. OAS1, OAS2, and OAS3 are part of the IFN-regulated double stranded RNA-activated antiviral pathway [73]. When OAS enzymes detect double stranded RNA, they synthesize 2’,5’-oligoadenylates, which then activate RNase L to degrade viral RNA leading to apoptosis and inhibition of virus replication [74–77]. Notably, other coronaviruses besides SARS-CoV-2 have been shown to produce viral proteins that target the degradation of OAS-RNase L pathway proteins, in order to reduce RNase-L activity and inactivate the host defence [78, 79]. OASL has also been suggested to play a role in enhancing antiviral innate immunity [80]. Thus, therapies that can enhance the levels and action of these anti-viral mediators bear a potential for the prevention of SARS-CoV-2 transmission.

Our results demonstrating increased apoptosis in cells treated with CBD and transfected with SARS-CoV-2 viral genes suggests a potential protective effect of CBD at initial infection. However, it also raises the question of whether this could be harmful in an individual who already had a high viral load. Currently, limited information is available on the use of CBD in patients with COVID-19. Based on the anti-inflammatory effects of CBD on the acquired immune system, there have been calls for the use of CBD in COVID-19 patients to treat acute respiratory distress syndrome (ARDS) [81], and to reduce the viral load [82]. In a murine model of ARDS, CBD administration downregulated levels of proinflammatory cytokines, and ameliorated clinical symptoms [83]. There is medical interest in the use of CBD to treat advanced SARS-CoV-2 infections, with eight clinical trials currently underway [84], including one studying use of CBD treatment for severe and critical COVID-19 pulmonary infection [85]. One trial has recently reported results, indicating no significant effect of 300 mg CBD daily on the clinical evolution of COVID-19 in patients presenting with mild to moderate symptoms, although the authors suggested that future studies should evaluate higher doses, as well as the clinical efficacy of CBD in patients with more severe COVID-19 [86]. Although results have not yet been reported from most other registered clinical trials, none have been stopped prematurely by the medical oversight committees, indicating that findings of significant harm have not been detected. It is therefore possible that CBD may offer prophylaxis against initial viral infection through a pro-apoptotic mechanism that does not result in widespread cell death in highly infected patients. Additional work will be required to understand the nature of CBD effects, in this regard.

## 5. Conclusions

Taken together, our results indicate that while expression of the SARS-CoV-2 genes *ORF8*, *ORF10*, and *M protein* alone fails to significantly induce apoptosis, or reduce cell numbers, and while treatment of cells with up to 2 μM CBD also does not affect these parameters, combinations of 2 μM CBD with these genes dramatically upregulates apoptosis and reduces cell numbers. A poor ability of cells to sense and respond to the presence of these viral genes may therefore be a factor in the high infectivity rate of SARS-CoV-2. The induction of Type II and Type III *IFN*, as well as *OAS* family member genes, may help explain the pro-apoptotic effect of CBD that was observed in cells expressing viral genes, and future work should investigate a causal role. In addition, the induction of these *IFN* and *ISG* by CBD in control cells may indicate a ‘priming’ effect on the innate immune system, better readying cells to respond to viral infection, which could help to explain the lower rates of COVID-19 in patients receiving high-dose CBD treatment.

## Acknowledgements

This work was supported by grants from the Natural Sciences and Engineering Research Council of Canada (NSERC) to R.E.D. #RGPIN-2019-05642 and RGPAS-2019-00008, and the Canada Foundation for Innovation—Leader’s Opportunity Fund and Ontario Research Fund (Project No. 30259). MFF was supported by a Mitacs COVID-19 Accelerate Postdoctoral Fellowship. The funders had no role in study design, data collection and analysis, decision to publish, or preparation of the manuscript.

